# Assessing the defensive behaviour of *Apis mellifera scutellata* in its native range reveals its variability, organization and persistence

**DOI:** 10.64898/2026.09.30.755645

**Authors:** Daniela Ramirez-Moreno, Max Fritzler, Ann-Kathrin Pohle, Kavitha Kannan, Emily Lippman, James C. Nieh, Kevin Balkwill, Morgane Nouvian

## Abstract

Colony defence is critical for the survival of wild honeybees but presents a challenge for local populations and beekeepers. *Apis mellifera scutellata* is an important pollinator in its native range, a large area of Southern and Eastern Africa ^1^, but is reputed for its explosive temperament. Here, we present data demonstrating that this behaviour is only exhibited by a small fraction of colonies, while others are relatively docile. This sheds new light on this subspecies and on its likely evolutionary trajectory. We further characterize the different elements of this defensive response and compare them to that of *A. m. carnica* and *A. m. mellifera*. We show that each subspecies invests differently in the two steps of the behavioural sequence (alert and attack). Furthermore, our results suggest that the task of attacking is carried out by specialized workers in *carnica*, but not in the other subspecies tested, highlighting subtle differences in colony organization across subspecies. Finally, we discovered that defenders from *A. m. scutellata* and *scutellata*-hybrid colonies can locate a target that was stung up to 2 h and 3 h earlier, respectively, whereas European bees could only do so for 30 min. This long-term marking could confer an advantage against returning predators with few weak points, such as honey badgers and humans wearing protective clothing. Overall, we offer a fresh perspective on this controversial but important subspecies of honeybees, with implications for both future studies and daily management.

## Results and discussion

Invasive species typically have reduced genetic variation because they derive from a small number of displaced individuals ^2,3^. This founder effect can fix particular phenotypes in the invasive population, misrepresenting the true characteristics of a species in its native range. The East African lowland honeybee, *Apis mellifera scutellata*, is infamous for giving rise to the highly aggressive “Africanized bees” - sometimes referred to as “killer bees” - in America. The aggressive phenotype is a dominant trait ^4–6^ inherited from a few colonies of *scutellata* that were imported to South America and crossed with European bees, in an attempt to breed a race of honeybees better suited to the tropics ^7^. While the defensive behaviour of these Africanized hybrids has been extensively characterized ^4,8–10^, few studies have been conducted on natural populations of *scutellata* in Africa. They are generally considered to be highly defensive, like their hybrid descendants, although some evidence contradicts this assumption. Indeed, the few studies ^11–13^ that did test the defensive behaviour of *scutellata* in its native range present a much more nuanced view: for example, in one study, only 2 out of 71 colonies exhibited violent responses such as the ones attributed to Africanized bees ^12^.

### The defensive behaviour of *A. m. scutellata* is highly variable

In February 2024, we surveyed defensive responses in 125 colonies of *A. m. scutellata* housed in a single apiary near Pullen Nature Reserve, a field station of the University of the Witwatersrand (South Africa). Each colony was fitted with an extended landing board and then a black leather flag was moved up and down in front of one entrance by a small motor for 2 min (Fig. 1A). We recorded the reaction time (time elapsed before the first sting), counted the number of stingers embedded in the flag at the end of the assay and rated pursuing efforts on a subjective scale from 0 (no bees followed the experimenter) to 5 (the experimenter had to leave the apiary). Each colony was tested only once.

**Figure 1:**
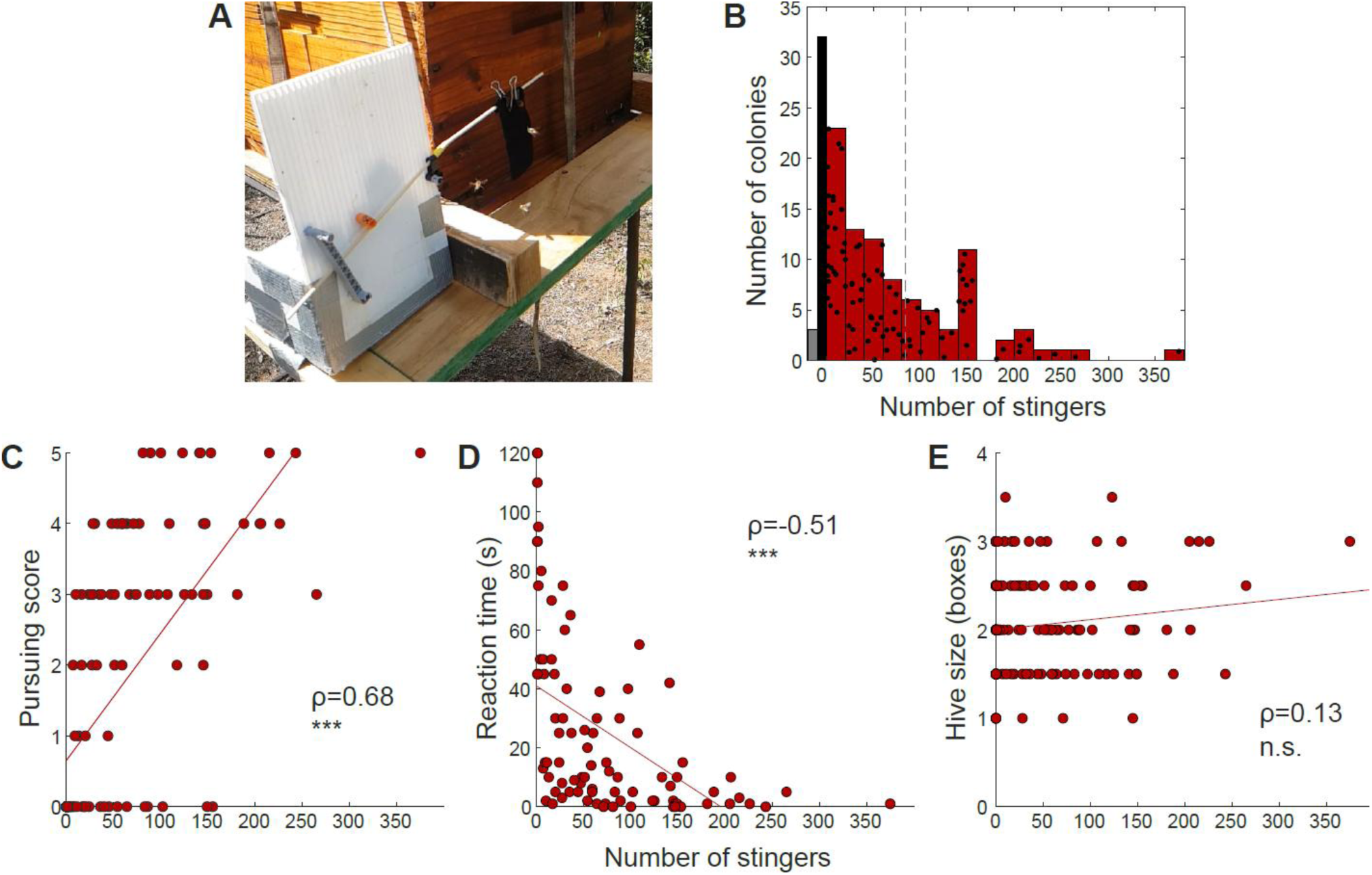
Colony defensiveness in *Apis mellifera scutellata.* A. Photo of the flag assay. A black leather flag was jiggled ∼7 cm from the entrance by a small motor. The total number of stingers embedded in the flag after 2 min was then counted. B. Distribution of the stinging responses. Each black dot is a unique colony, the red bars summarize the number of colonies with responses within each bin of 20 stingers. The black bar sums the 32 colonies that did not respond (no stinger in the flag) while the grey bar denotes the 3 colonies that retreated inside the hive instead of attacking. In total, 125 colonies were tested. For comparison, the dashed line indicates the maximum number of stingers measured in European bees (*Apis mellifera mellifera)* with a similar protocol. C. The subjective intensity of pursuit after the test is correlated with the number of stingers delivered to the flag (Pearson, n=125). D. The reaction time (delay between the start of the disturbance and the first stinging event, when some bees did sting) is negatively correlated with the number of stingers delivered (Pearson, n=86). E. The size of the colony, estimated by the number of hive boxes (with 1 for the main body and 0.5 for each honey super) is not correlated with the number of stingers delivered (Pearson, n=125).

The range of stinging responses we observed was incredibly wide (Fig. 1B). In about a quarter of colonies (35), no bees stung the flag at all. Of these, 3 colonies even retreated inside the hive such that the landing board was empty and all traffic had ceased by the end of the assay. At the other end of the spectrum, in 27 colonies more than 100 bees stung the flag, with a maximum at 375 stingers. Note that in these extreme events, the flag was covered by such a dense ball of bees that new defenders could not reach the leather anymore, so this count is likely underestimating the number of attacking bees. Given that all colonies were housed in a single location and were tested within 8 days of relatively stable weather conditions, this variability in defensive behaviour among *A. m. scutellata* colonies likely represents true genetic diversity, confirming previous works ^11–13^. Most studies on Africanized bees report stinging rates above 50 stings per minute (see the comprehensive overview provided in Table S6 of ^14^, and references therein), although direct comparisons are complicated by methodological variability (in particular regarding the use of a prior disturbance and the location of the target presented) and possible seasonal effects. This high defensiveness was attributed back to *scutellata* and interpreted as an adaptation to high predation rates ^15,16^. Our results challenge this view and suggest that Africanized hybrids are derived from a set of colonies that were among the most defensive in native *scutellata* populations. As a consequence, an alternative interpretation of the evolutionary trajectory of this defensive behaviour, already proposed by Rinderer ^7^, seems more likely: that *scutellata* retained this diversity because honey harvesting from this subspecies relies on non-selective trapping (and until recently, destruction) of wild swarms, while beekeeping practices have selected against high aggressiveness in European bees. At the root of these different handling methods is the tendency of *scutellata* colonies to abscond easily, hence making colonies hard to keep for a long time, whereas European honeybees establish perennial colonies more amenable to human management.

The number of stinging bees was positively correlated with pursuing behaviour (Fig. 1C, Pearson’s, ρ=0.68, p<0.001) and negatively correlated with reaction time (Fig. 1D, Pearson’s, ρ=-0.51, p<0.001). Reactions were usually fast, with half (44 out of 86) of responsive colonies stinging within 10 s of the flag starting to move, but again there was high variability, with 12 colonies taking more than 60 s to initiate defence. This is consistent with the observations made in Africanized bees ^8,14^. The size of the colony, estimated by the number of hive boxes, did not appear to correlate with the intensity of the stinging attack (Fig. 1E, Pearson’s, ρ=0.13, p=0.137). A more direct measure of colony size would have been needed to confirm this observation, which is at odds with a previous report ^12^.

### Time course of the defensive response of *A. m. scutellata*

In order to further characterize the defensive response of *scutellata,* in January 2025 we repeatedly tested 3 colonies established at Pullen Nature Reserve. For each trial, a white billboard was placed on the side of the hive to provide a background on which flying bees could be counted (Fig. 2A). A first camera recorded the activity on the landing board (top view) and a second on this billboard (side view). Baseline activity was recorded for 5 min, then the flag assay was performed for 2 min as before. We kept recording for 8 minutes afterwards, with the flag removed, to observe the recovery. One frame was extracted from the recordings every minute, and the number of bees seen flying in front of the billboard (Fig. 2B, upper panel) or standing on the landing board (Fig. 2B, lower panel) was counted. The data was then normalized by subtracting the baseline (average of timepoints-5 to 0 minutes).

**Figure 2:**
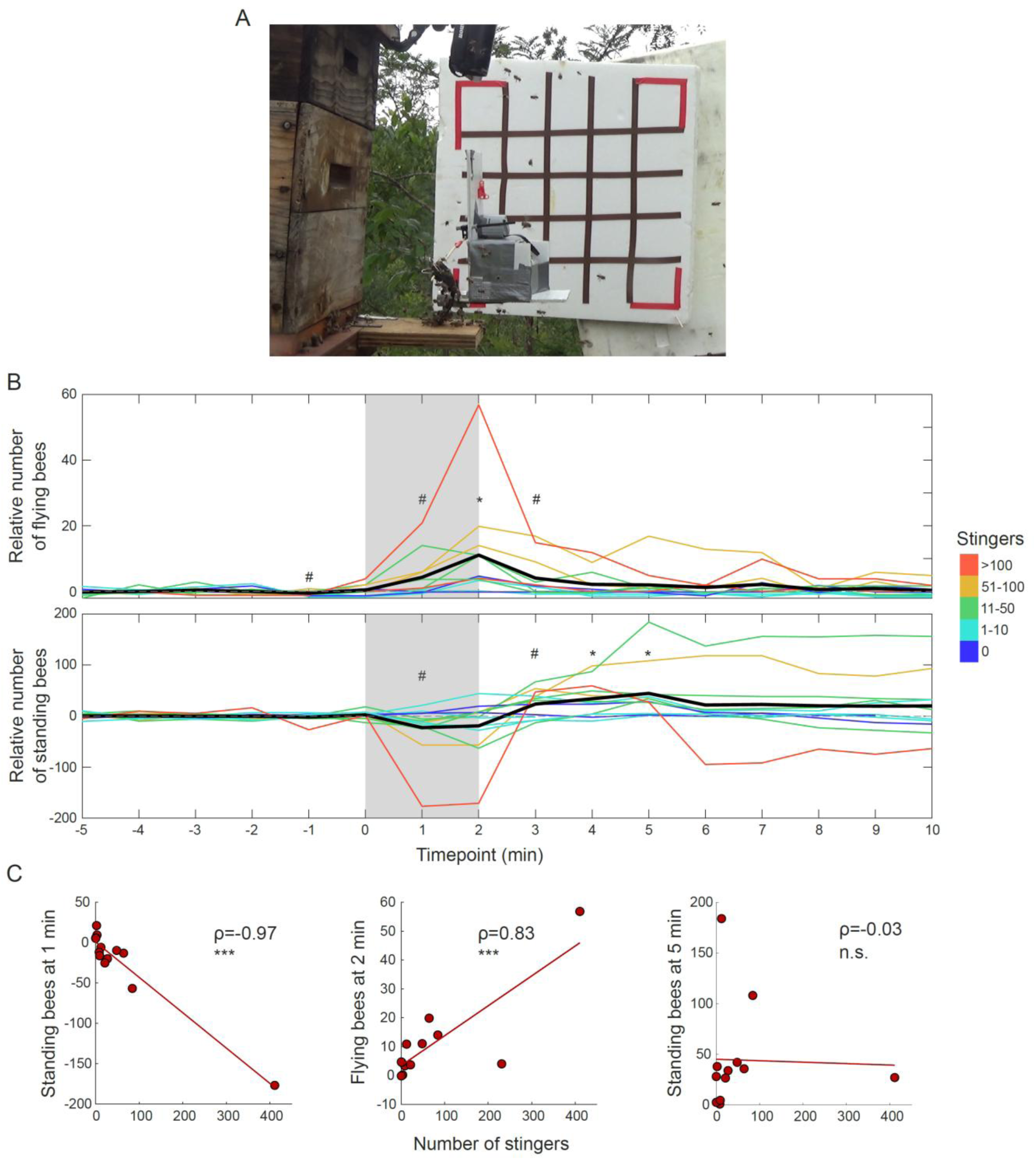
Timecourse of the stinging attack. A. Photo of the assay (from the lateral camera), showing the billboard, the flag and the second camera looking down on the landing board. B. Number of flying (upper panel, n=12 trials) and standing (lower panel, n=14 trials) bees at each timepoint, relative to the average number of bees performing each behaviour before the disturbance. * highlights significant differences from baseline, # shows timepoints which were different before FDR correction. Thin lines are single trials, colour-coded according to the number of stingers delivered to the flag; the thick black line is the average of all trials (N=3 colonies). C. The total number of stingers delivered to the flag is correlated with a dip in the number of standing bees 1 min after the start of the disturbance and with an increase in flying bees after 2 min, but not with the number of standing bees on the landing board after 5 min (Pearson).

During the disturbance, the number of bees on the landing board tended to decrease, albeit this was not significant after correcting for multiple comparisons (Wilcoxon signed rank test, timepoint +1 min: p=0.048 before FDR correction, p=0.191 after). This is most likely because bees moved from the landing board to the flag, and indeed the depth of this dip in the number of bees on the landing board was correlated with the number of stingers in the flag (Fig. 2C, left panel, Pearson’s, ρ=-0.97, p<0.001). In addition, the number of flying bees increased progressively, reaching a peak at the end of the disturbance (Wilcoxon signed rank test, timepoint +2 min: corrected p=0.023). This increase was correlated with the intensity of the attack as measured by the resulting number of stingers in the flag (Fig. 2C, middle panel, Pearson’s, ρ=0.83, p<0.001). As experimenters, we can attest that this flying activity is a very striking feature of the defensive response of *scutellata* that we never observed in such proportion in European subspecies. It likely serves to threaten predators while preserving the life of the defenders, unlike the more efficient but suicidal act of stinging. After the disturbance, the number of flying bees returned to baseline quickly but the number of bees on the landing board became higher than baseline for several minutes (Wilcoxon signed rank test, timepoint +4 min: corrected p=0.039, timepoint +5 min: corrected p=0.008). Interestingly, this increase was not correlated with the stinging response (Fig. 2C, right panel, Pearson’s, ρ=-0.03, p=0.923), nor with flying activity before (Pearson’s, ρ=-0.08, p=0.843) or during (Pearson’s, ρ=0.02, p=0.956) the disturbance. Thus, this higher number of bees on the landing board likely represents an increase in guarding effort in case the predator returns, rather than the consequence of returning foragers or defenders landing after the disturbance.

### Comparisons with *A. m. carnica* and *A. m. mellifera*

In the summer of 2021, we conducted experiments using a similar disturbance protocol on 5 colonies of *A. m. carnica* (Carniolan bees) and 7 colonies of *A. m. mellifera* (European dark bees) hosted respectively at the University of Konstanz (Germany) and in Tägerwilen (Switzerland), about 6 km apart. We took advantage of this existing dataset to compare the defensive behaviour of these European subspecies to that of *scutellata.* Note that the phylogenetic relationships between these subspecies have been challenging to uncover ^17–21^. All studies agree that *carnica* and *mellifera,* despite their close geographical proximity, derive from independent migrations and thus belong to separate lineages. *A. m. mellifera* sometimes appear more closely related to *scutellata* than to *carnica,* which has led to the hypothesis that they entered Europe from Africa, through the Iberian peninsula ^17,19,21^. However, other studies, including the most recent and comprehensive one, argue for an Asian origin of *Apis mellifera,* which clearly sets the African lineage aside from the European ones ^18,20^.

Recruitment via sting alarm pheromone (SAP) release is a crucial element of the defensive behaviour of honeybees ^22,23^. This can happen through the accumulation of stingers in the target, but also through a specific behaviour in which a bee raises its abdomen and extrudes its stinger. We quantified the number of bees performing this active release of the SAP during the disturbance in each of the 3 subspecies and found striking differences (Fig. 3A). This behaviour was rarely seen in *carnica* but frequent in *mellifera* and *scutellata* (GLMM, main effect of subspecies: Χ^2^=13.282, p=0.001, post-hoc pairwise comparisons corrected with FDR: *carnica* vs *mellifera*: p<0.001, *carnica* vs *scutellata*: p<0.001, *mellifera* vs *scutellata*: p=0.112). It has been proposed that the high defensiveness of Africanized bees is caused in part by increased responses to the SAP ^24^. Our results point to an alternative (non-exclusive) mechanism, namely that some subspecies release higher concentrations of SAP to start with. Although the end result is similar (stronger colony defence), it is worth noting that these 2 mechanisms rely on different stimuli (olfactory vs visual) and different types of bees (recruits vs guards), which have important consequences for subsequent studies.

**Figure 3:**
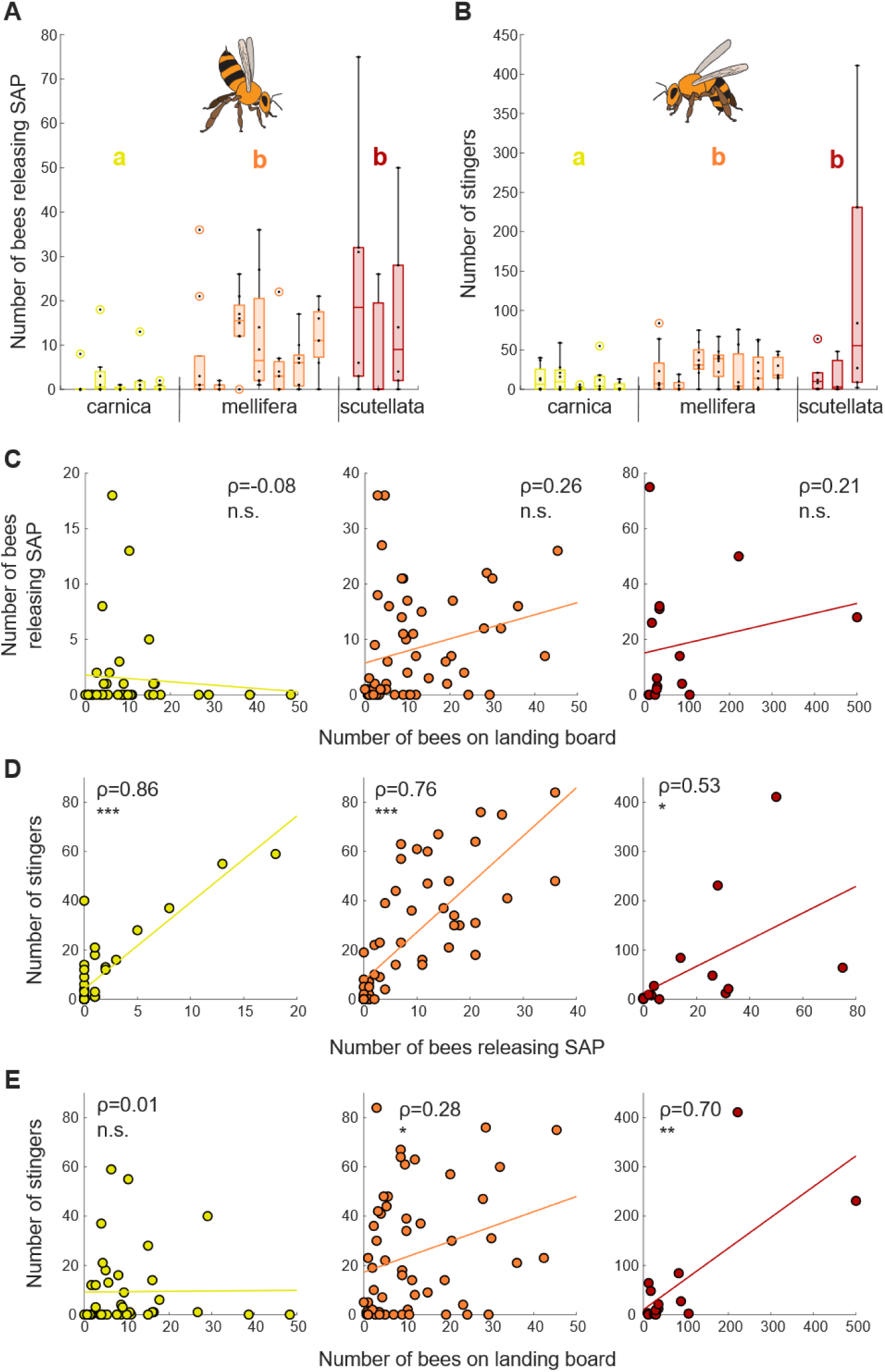
Comparisons between A. m. carnica, A. m. mellifera and A. m. scutellata. A. Number of bees actively releasing the sting alarm pheromone during the 2 min of disturbance, for colonies of the 3 subspecies tested repeatedly. Each boxplot is the data from a single colony, each black dot is a single trial. Different letters indicate significant differences at the subspecies level. B. Number of stingers in the flag after 2 min, for the same colonies as in panel A. Each boxplot is the data from a single colony, each black dot is a single trial. Different letters indicate significant differences at the subspecies level. C. The number of bees visible on the landing board before the disturbance is not correlated with the number of bees actively releasing the sting alarm pheromone in any subspecies. D. The number of bees actively releasing the sting alarm pheromone is correlated with the number of stingers delivered to the flag in all subspecies. E. The number of bees visible on the landing board before the disturbance is correlated with the number of stingers delivered to the flag in *A. m. mellifera* and *A. m. scutellata*, but not in *A. m. carnica.* carnica: N=5 colonies, n=38 trials (7-8 per colony); mellifera: N=7 colonies, n=57 trials (7-9 per colony); scutellata: N=3 colonies, n=15 trials (3-6 per colony).

The number of stingers embedded in the flag at the end of the 2 min of disturbance was also linked to the subspecies considered (Fig. 3B). Overall, *carnica* was less defensive than either *mellifera* or *scutellata* (GLMM, main effect of subspecies: Χ^2^=7.155, p=0.028, post-hoc pairwise comparisons corrected with FDR: *carnica* vs *mellifera*: p=0.027, *carnica* vs *scutellata*: p=0.018), while the comparison between *mellifera* and *scutellata* was not significant (p=0.360). However, generalizing about *scutellata* defensive behaviour is difficult since, as before, the variability between *scutellata* colonies was very high, with one colony clearly presenting extreme responses (last colony on Fig.3B). Note that the variability between trials for this colony is also important, with responses ranging from only 2 stingers on the flag to a record of 411 stingers. It is also worth keeping in mind the unavoidable differences in environmental conditions due to *scutellata* not being sympatric with *carnica* and *mellifera*, that could impact the expression of defensive behaviour.

Interestingly, the number of bees actively releasing SAP was not correlated with the number of bees on the landing board at the start of the disturbance in any of the 3 subspecies (Fig.3C, Pearson’s, *carnica*: ρ=-0.08, p=0.615, *mellifera*: ρ=0.26, p=0.060, *scutellata*: ρ=0.21, p=0.461), indicating separate workforces for foraging and alerting. However, consistent with the proposed recruiting function of this behaviour, active SAP release was correlated with the stinging response (Fig.3D, Pearson’s, *carnica*: ρ=0.86, p<0.001, *mellifera*: ρ=0.76, p<0.001, *scutellata*: ρ=0.53, p=0.042). We could expect the number of stingers to also be correlated with the number of bees on the landing board at the start of the disturbance, as these would be potential recruits. This is indeed the case for *mellifera* and *scutellata* but not for *carnica* (Fig. 3E, Pearson’s, *carnica*: ρ=0.01, p=0.956, *mellifera*: ρ=0.28, p=0.037, *scutellata*: ρ=0.70, p=0.003). This suggests that *carnica* colonies may need to recruit a specialist workforce from inside the colony to defend, whereas defence and foraging may overlap in *mellifera* and *scutellata*, facilitating recruitment and (in combination with the higher SAP release described above) ultimately generating stronger defensive responses. Such a difference in organization between European and Africanized bees was hypothesized by Breed et al. after they characterized “soldier bees” in colonies of European descent ^25^. Although they did not specify the specific subspecies that they used in this study, their colonies likely belonged to the C lineage (which includes *carnica* and *ligustica*) as it is the most common in the USA ^20^.

### Predator marking

While surveying the defensive behaviour of *scutellata* colonies, we noticed that bees would keep attacking a stung area long after the stingers had been removed. We suspected that a substance remained on the stung material and marked it. To test this hypothesis, we collected stung flags by disturbing donor colonies, removed the stingers, and stored the flags individually for a defined period (from 5 to 300 min). We then presented a previously stung flag along with a clean, control flag to a test colony while disturbing it, and allowed both flags to collect stingers (Fig. 4A). This experiment was conducted (with small variations in protocols detailed in the Methods) on 3 populations of bees: *scutellata* in South Africa, European bees in Germany, and *scutellata*-hybrids in the USA. In all three datasets, the time elapsed since the treated flag was stung significantly affected the proportion of stingers that it collected during the test (GLMM, *scutellata* bees: *F*_4,63_ = 11.82, *P* < 0.0001; European bees: *F*_3,95_ = 11.66, *P* < 0.0001; hybrid bees: *F*_4,33_ = 3.75, *P* = 0.0127). When doing this procedure with *scutellata* colonies, bees still focused their attack preferentially on the previously stung flag up to 120 min after the initial attack (Fig. 4B, predicted fraction of stingers directed to the previously stung flag with a model-weighted mean initial loading of 25.9 stingers at 30 min: 0.900, 95% CI = 0.837–0.941, at 60 min: 0.796, 95% CI = 0.710–0.861 and at 120 min: 0.660, 95% CI = 0.544–0.760). After 180 min, both flags were stung in statistically indistinguishable proportions, suggesting that bees could no longer detect any marking substance on the previously stung flag (predicted fraction of stingers on previously stung flag at 180 min: 0.578, 95% CI = 0.474–0.676, at 300 min: 0.604, 95% CI = 0.455–0.736).

**Figure 4:**
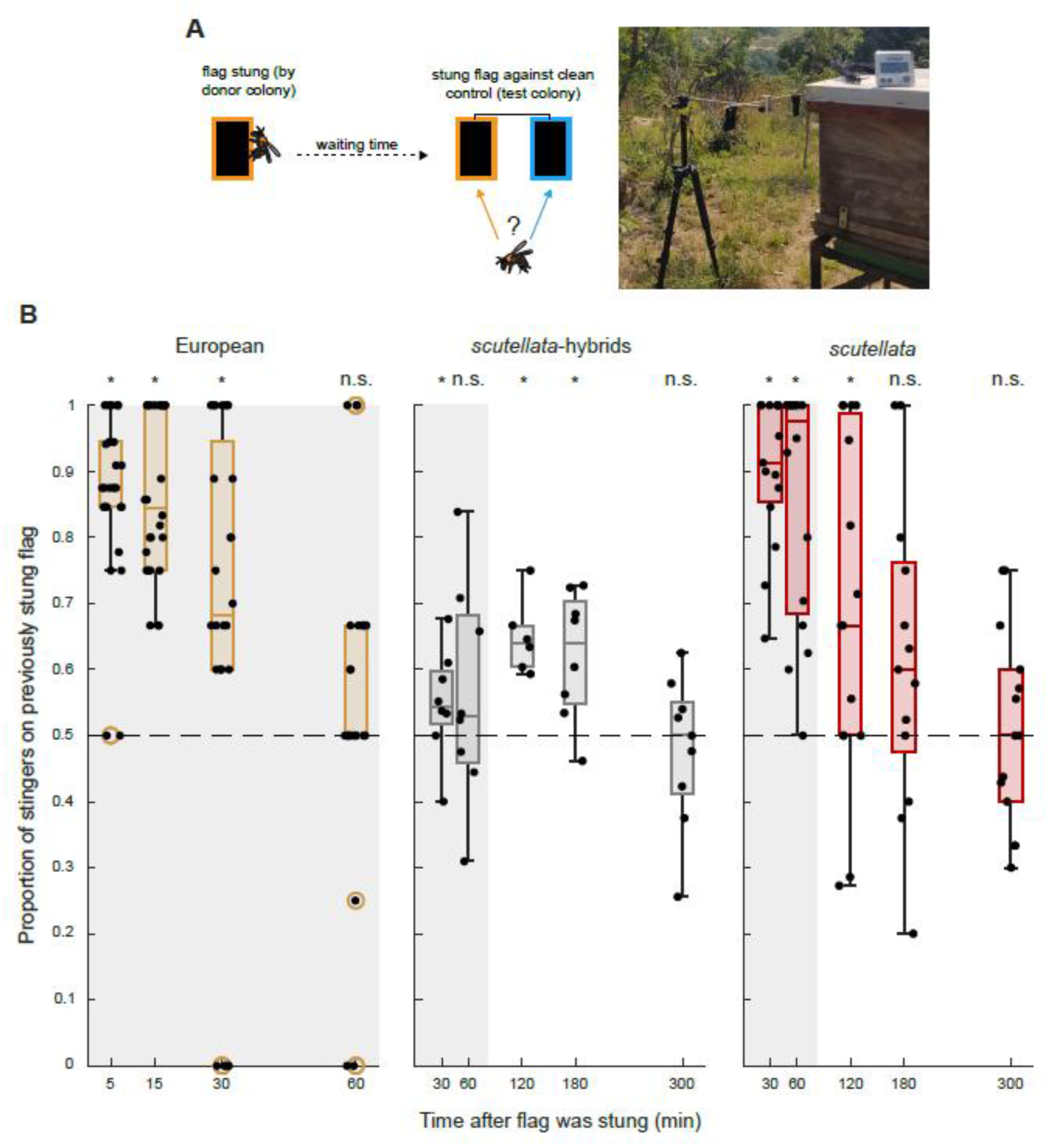
Marking of enemies in European bees, *scutellata*-hybrids and *scutellata.* A. Experimental protocol. A leather flag was waved in front of a donor colony to collect stingers. The stingers were then removed, and the flag stored for a defined duration (from 5 to 300 min). After this time, the stung flag was presented to another colony along with a clean flag, both equidistant from the colony entrance. The colony was disturbed until they started stinging the flags, at which point the attack was allowed to continue for a few minutes. The number of stingers in each flag was then counted. B. Proportion of stingers on previously stung flag (0: all stingers on control flag, 0.5: equal number of stingers on both flags, 1: all stingers on previously stung flag) as a function of the delay between the two attacks. European bees (brown) still targeted the flag that was already stung 30 min after the first attack but did not differentiate between the two flags after 60 min. *scutellata*-hybrids (grey) and *scutellata* (red) targeted the stung flag for much longer, up to 180 and 120 min respectively. Note the difference in scale of the x-axis, such that the shaded area represents the same duration (60 min) in all graphs. The black dots correspond to single trials. European: N=12 colonies, n=100 trials; *scutellata*-hybrids: N=8 colonies, n=39 trials; *scutellata*: N=6 colonies, n=73 trials.

The *scutellata*-hybrids exhibited a very similar pattern: at a model-weighted mean initial loading of 44.72 stingers, the predicted fraction of stingers on the previously stung flag was above chance level for up to 180 min (model predictions at 30 min: 0.566, 95% CI = 0.501–0.629, at 120 min: 0.636, 95% CI = 0.563–0.703 and at 180 min: 0.587, 95% CI = 0.526–0.645). The 60 min timepoint was not significant but the confidence interval overlapped only slightly with chance level (0.558, 95% CI = 0.493–0.621), thus it is likely due to low statistical power and biological noise rather than to a true temporary loss of the marking effect. As before, marking was no longer detectable at 300 min (0.493, 95% CI = 0.434–0.552). The longer persistence found in these hybrids compared to pure *scutellata* may be driven by the heavier initial stinger load on the treated flag (41.1 stingers on average for hybrids, compared to 24.5 for *scutellata*). Alternatively, the production or perception of the marking compound(s) could have been impacted by a founder effect in the invasive hybrids, resulting in a more persistent phenotype.

Finally, we repeated this experiment with European colonies in Konstanz (Germany), after adjusting the waiting periods based on pilot data. We found that the previously stung flag collected more stingers only up to 30 min after the initial attack (Fig. 4B, predicted fraction of stingers directed to the previously stung flag at the mean initial loading of 15.11 stingers at 5 min: 0.865, 95% CI = 0.817–0.902, at 15 min: 0.860, 95% CI = 0.798–0.906, and at 30 min: 0.724, 95% CI = 0.621–0.807). Marking was no longer detectable after 60 min (0.558, 95% CI = 0.427–0.681). Although the initial stinger load was lower in this experiment (only 15.1 stingers on average), which could have driven the apparent shorter persistence of the marking in this subspecies, repeating the 60 min timepoint with a much higher loading (33.67 stingers on average) did not change the result (Suppl. Fig. 3, Wilcoxon signed rank test against chance level, p=0.469).

The main vertebrate predators of African honeybee colonies are humans and honey badgers (*Mellivora capensis*) ^11,26^. Both pose a similar challenge to defending bees: humans regularly wear protective clothing, and honey badgers have notoriously tough skin ^27^, making them hard to sting. In addition, honey badgers have been reported to return multiple times to the same colony over days, until the brood is entirely consumed ^28^. Hence, marking the few areas that are vulnerable to stings could provide critical information to honeybees for deterring these predators. The origin and nature of this marking substance is unknown, but it seems logical that it would be part of the SAP or of the venom. We could not find data on the composition of either of these secretions in *scutellata*, but there is some information from Africanized hybrids. Comparisons of their SAP with that of European subspecies identified a single compound, 3-methyl-2-buten-1-yl acetate, as being produced exclusively in Africanized honeybees ^29^. However, the high volatility of this compound (vapor pressure of 3.99 mmHg at 25°C) makes it an unlikely candidate for such long-term marking. A more convincing candidate could be octyl acetate: it has lower volatility (0.19 mmHg at 25°C), provides information about target location to stinging bees ^30^, and is produced in much higher quantities in Africanized honeybees ^31^. Alternatively, the marking compound(s) could be part of the venom, which would explain why only the areas that were successfully stung appeared marked (painful personal observations). The venoms of Africanized and European bees are complex and broadly similar, but still noticeably different ^32–34^.

## Conclusions

Our study highlights a number of overlooked features in the defensive behaviour of *Apis mellifera scutellata*, consequently shedding new light on how this behaviour may have evolved and providing crucial information for future studies on this important behaviour. More precisely, we demonstrate that this behaviour is highly variable and likely organized differently than the one of the more popular *carnica* subspecies, including more SAP release by guards, generalist recruits and flying defenders. We also discovered the existence of a long-term marking of stung areas, critical information for beekeepers handling this unpredictable subspecies.

## Methods

### Survey of colony defence in *scutellata*

A total of 125 colonies were assessed for their defensive responses. They were located in a single apiary in Moederlief (Mbombela, South Africa; 25°32’25.1“S, 31°12’26.1”E). All colonies were wild caught around either Johannesburg or Mbombela, where *scutellata* is the only established subspecies (occasional occurrences of *capensis* are systematically destroyed by beekeepers, as they provoke colony collapse by parasitizing *scutellata* colonies). The survey was conducted over 8 days in February 2024, corresponding to high summer in this area. A trial went as follows. First, we checked that the colony was active (by the regular shuttling of foragers) and fitted it with a 15 cm wide landing board, taking care to disturb the bees as little as possible. This was done to facilitate observation and to standardize the testing conditions, as some colonies had small landing boards, whereas others had none. We then waited 5 min to make sure that the colonies were calm. A black suede leather flag (10 x 5 cm), mounted on a small motor (Lego Power Functions XL-Motor), was then placed about 7 cm from the colony entrance (Fig. 1A). This flag was jiggled for 2 min, with no other form of disturbance, and then quickly placed in a plastic box to stop the attack. We noted: 1) the size of the hive as the number of boxes it was made of, counting the main hive body as 1 and any additional honey supers as 0.5 since they are smaller, 2) the reaction time, i.e. the delay between switching on the motor to start the flag movement and the first sting received by the flag, 3) the number of stingers embedded in the flag after the 2 min of stimulation and 4) the pursuing effort as a subjective measure of the number and persistency of the bees following the experimenter after the trial, on a scale from 0 (no pursuers) to 5 (the experimenter had to leave the apiary). Importantly, we did not start the next trial until all pursuers were gone and we always moved to a distant colony, to avoid cross-reactions.

### Detailed characterization of the defensive response in *scutellata*

Additional information about how the defensive response of *scutellata* colonies unfolds was obtained by disturbing 3 colonies located in Pullen Nature Reserve (University of the Witswatersrand, South Africa, 25°34’26.6“S, 31°10’58.9”E) in January 2025. A square landing board of roughly 17 x 17 cm was fixed permanently in front of the colony entrance and a billboard with a grid was placed on the side of the colony at entrance level (Fig. 2A). They provided standard areas on which to count walking and flying bees, respectively. Two cameras (Canon Legria and Sony HDR-CX625) were used to record the trial, one fixed to the top of the colony and pointing down on the landing board, the other on a tripod and aimed at the billboard. We started the video recordings 5 min before the start of the disturbance and stopped them 10 minutes after. The disturbance itself was the same as before, i.e. a black flag jiggled for 2 min in front of the entrance. In total we conducted 15 trials (colony 1: n=6, colony 2: n=3, colony 3: n=6).

Because of diverse issues, 3 videos of the billboard could not be used at all, and 3 videos of the landing board (from different trials) were cut short (total durations of 5 min, 8 min, and 10 min). This truncated data was included nonetheless. To analyse the videos, frames were automatically extracted at 1 min intervals after setting the start of the disturbance as reference (timepoint 0). If a bee happened to be on the camera lens and blocking the view, another frame was chosen as close as possible to the given timepoint. The number of bees seen flying in front of the billboard or standing on the landing board was manually counted in each frame. During one trial, the landing board was covered with over 500 bees at the start of the disturbance, making it impossible to count them accurately. We included this estimate as a measure of the number of bees on the landing board before the disturbance (Fig. 3C & E) but excluded it from the time course analysis (Fig. 2B).

Finally, we also used the videos of the landing board to count the number of bees actively releasing the SAP during the 2 min of disturbance. This behaviour was defined as the bee lifting its abdomen to point it upwards (see schematic in Fig. 3A), an unusual stance that also caused a characteristic walking pattern. To perform this scoring, the landing board was subdivided into small squares by placing a grid on the computer screen and each square was carefully observed during the 2 min of the disturbance. If a bee was seen to move across multiple squares while performing this behaviour, it was noted to avoid counting the same bout twice.

### *Carnica* and *mellifera* dataset

The 5 colonies of *carnica* were located at the University of Konstanz (Germany; 47°41’ 27.5“N, 9°11’17.1”E), and the 7 colonies of *mellifera* were 5.7 km away in Tägerwilen (Switzerland; 47°39’36.6“N, 9°07’33.3”E), in an apiary dedicated to the breeding of this subspecies and part of the “mellifera.ch” association, which undergoes regular genetic tests. Since both *carnica* and *mellifera* are common subspecies in the area, we double checked the genetics of the tested colonies in 2 ways. We collected an antenna from 30 drone pupae in each of our *carnica* colonies and sent them to Apigenix, an Institute for bee genetics, for sequencing. Based on single nucleotide polymorphisms ^35^, all colonies were assigned to the C lineage (i.e. *carnica* or *ligustica*) with more than 95% probability. We also used wing morphometrics ^36^. We sampled the left forewing of 15 workers per colony and photographed them under a binocular microscope (Leica S6D, camera Leica 10445930 1.0x). We then used the program TpsDig ^37^ (tpsDIG2w64, version 2.31) to place 18 landmarks on the vein junctions of each wing, following the protocol by Tofilski ^36^. A principal component analysis on the coordinates of these landmarks clearly separated the two subspecies (Suppl. Fig. 2; permutation test: p<0.001), as expected.

The experiments were conducted in July and August 2021, during summer. Each colony was tested 7 to 9 times, with at least 2 days between consecutive tests (final sample sizes: 38 trials for *carnica*, 57 for *mellifera*). We standardized the size and geometry of the hive entrances so that the number of bees responding to the stimulus would not be influenced by space constraints. All entrances were made into a rectangular slit, 1 cm in height and 22 cm in length, using white plastic boards hung on the hive front. Additionally, we attached a landing board (30 cm x 20 cm) made from the same material in front of the hive to facilitate observations. As for experiments with *scutellata*, care was taken to disturb the colonies as little as possible when placing these elements, and we waited 5 min before starting the test to ensure that they were calm. A camera (Panasonic, LUMIX GF 3) was placed on a tripod on the side of the colony entrance and recorded the activity on the landing board. We filmed the entrance area for 2 min without any disturbance to estimate baseline activity, then applied our disturbance protocol for 2 min. The motor used was the same as for *scutellata*, but the flag was slightly smaller (7 cm x 4 cm). In addition, the flag was placed at 4 cm from the entrance of the hive for the first minute and then shifted to a distance of 12 cm for the second minute. This was done because the main aim of this experiment was to quantify active SAP release. The flag being close to the entrance in the beginning favoured the initiation of the defensive response, while moving it further provided a better view of the bees performing this behaviour. For the later experiment with *scutellata*, we decided to simplify this protocol by choosing a constant, intermediate position for the flag.

### Predator marking experiment

This experiment is composed of 3 datasets, with small variations in protocol. The data on *scutellata* was collected using 6 colonies located in Pullen Nature Reserve. The data on European bees used 12 colonies located at the University of Konstanz. No genetic analysis was conducted on these colonies so they could be of *carnica, mellifera* or *ligustica* background. Finally, the data on hybrids was collected from 5 feral colonies at the Elliot Chaparral Reserve (San Diego County, California, USA) and from 3 colonies at the Biology Field Station (BFS) in La Jolla, CA on campus at UC San Diego. These colonies were known *scutellata*-hybrids based on prior genotype analysis ^14^ or maternal mitotyping ^38^. The test procedure consisted of 2 steps. The first step was to disturb a donor colony while also presenting a black leather flag (7 x 4 cm), in order to collect stingers on this flag. This disturbance was realized by waving a black ostrich feather in front of the colony for experiments with *scutellata* and European bees, or by dropping a brick (4 x 9.2 x 5.7 cm, 2 kg) 3 times on top of the colony, from a height of 30 cm, for hybrids. The flag received on average 24.5 ± 6.6 stingers in the *scutellata* dataset, 15.1 ± 0.7 stingers in the European dataset, and 41.1 ±19 stingers in the hybrid dataset. This flag was then quickly sealed in a plastic box to prevent additional attacks and brought back into the lab, where the stingers were counted and removed with forceps. It was stored for a given duration (5, 15, 30 or 60 min for experiments with European bees, 30, 60, 120, 180 or 300 min for experiments with *scutellata* and hybrids), either in an open plastic bag (*scutellata* and European bees) or hanging by tape from a desk (hybrids). The second step took place after the desired waiting period. For *scutellata* and European bees, the previously stung flag and a new, clean flag were mounted with paper clips on a small wooden pole at a distance of 30 cm from each other; for hybrids, the patches were mounted 20 cm apart on a rod. The pole or rod was placed at a distance of 30 cm, parallel to the front of a test colony (different from the donor colony) such that the flags were equidistant from the colony entrance (Fig. 4A). This colony was then disturbed as before (feather or brick), until the bees started stinging the flags. We allowed the flags to collect stingers for a short time (1 min in the *scutellata* and European datasets, 5 min in the hybrid dataset), at which point we quickly sealed them in plastic boxes again. Later, the number of stingers embedded in each flag was counted. The donor and test colonies were rotated, so that each colony contributed to both steps. Each colony contributed to all timepoints, but the sample sizes varied slightly. They are given in Suppl. Tables 1-3. Since the stinger load on the marked flag varied between datasets, it could have contributed to the effect. To check that the lower stinger load was not the reason for the shorter marking duration observed in European bees, an additional test was performed at the 60 min timepoint, on 6 new colonies, with an average initial load of 33.7 stingers. The results (Suppl. Fig. 3) confirmed that no marking could be observed after this duration.

## Statistics

The statistical analyses were conducted mainly in MATLAB R2024a. Correlations between variables were assessed using Pearson’s ρ. Variations in the number of flying or standing bees during the course of a defensive bout were tested using Wilcoxon’s signed rank test on each timepoint, and corrected for multiple testing with the Benjamini and Hochberg’s False Discovery Rate (FDR) procedure ^39^. To compare the SAP release and stinging behaviours of the 3 subspecies, we used a generalized linear mixed model with a Poisson distribution appropriate for counting data, with the subspecies as fixed effect and the colony identifier as random factor, followed by pairwise contrasts corrected with FDR. Finally, for each predator-marking dataset, we analysed the paired-choice response with a binomial generalized linear mixed model. Each assay presented one previously stung flag and one clean control flag simultaneously to the same test colony, so the model was specified conceptually as the number of stings on the previously stung flag out of the total number of stings on both flags. In JMP v19.1.10, this grouped binomial model was implemented as the fraction of total stings directed to the previously stung flag, weighted by the total number of stings in that assay, with a binomial error distribution and logit link. Because the sampling intervals and some protocol details differed among the *scutellata*, European, and hybrid datasets, we analysed them separately rather than pooling them in a single model. In each model, the delay between the 2 steps was treated as a categorical fixed effect, the original number of stingers loaded onto the previously stung flag was included as a covariate, and test colony was included as a random effect to account for repeated testing of colonies. The *scutellata* dataset used the full model, as both the initial stinger load (*F*_1,63_ = 4.29, p= 0.0425) and its interaction with the delay were significant (*F*_4,63_ = 3.18, *p* = 0.0192). By contrast, these terms were not significant in the other datasets (for European bees, stinger load: *F*_1,95_ = 0.16, p = 0.690, interaction: *F*_3,92_ = 0.16, p = 0.930; for *scutellata-*hybrids, stinger load: *F*_1,33_ = 0.46, p = 0.504, interaction: *F*_4,29_ = 1.77, p = 0.162), so the models were reduced by removing the interaction. To determine how long marking remained detectable, we used planned model-based estimates at the mean original stinger loading to test whether the predicted fraction of stingers directed to the previously stung flag exceeded the null expectation of 0.50. A time point was considered to show detectable marking when the 95% confidence interval for the predicted fraction was entirely above 0.50. These were planned persistence tests at the sampled ages, not post hoc all-pairwise comparisons among ages.

## Acknowledgements

We are very thankful to Feri Jusko, Jan Larsson and Maria Larsson for allowing us to bother their *mellifera* colonies. We also thank Gabriele Soland at Apigenix for her help with analysing the genetic background of our colonies. Marcus Byrne, Maleti Mpila, James Foster and Frida Hildebrand helped us with the logistics of our stay at Pullen and Promise Skhosana provided invaluable assistance for dealing with *scutellata*. We would also like to thank Jerry Zhiyuan and Zachary George for their help during experiments on *scutellata*-hybrids.

---

**Supplementary Figure 1:**
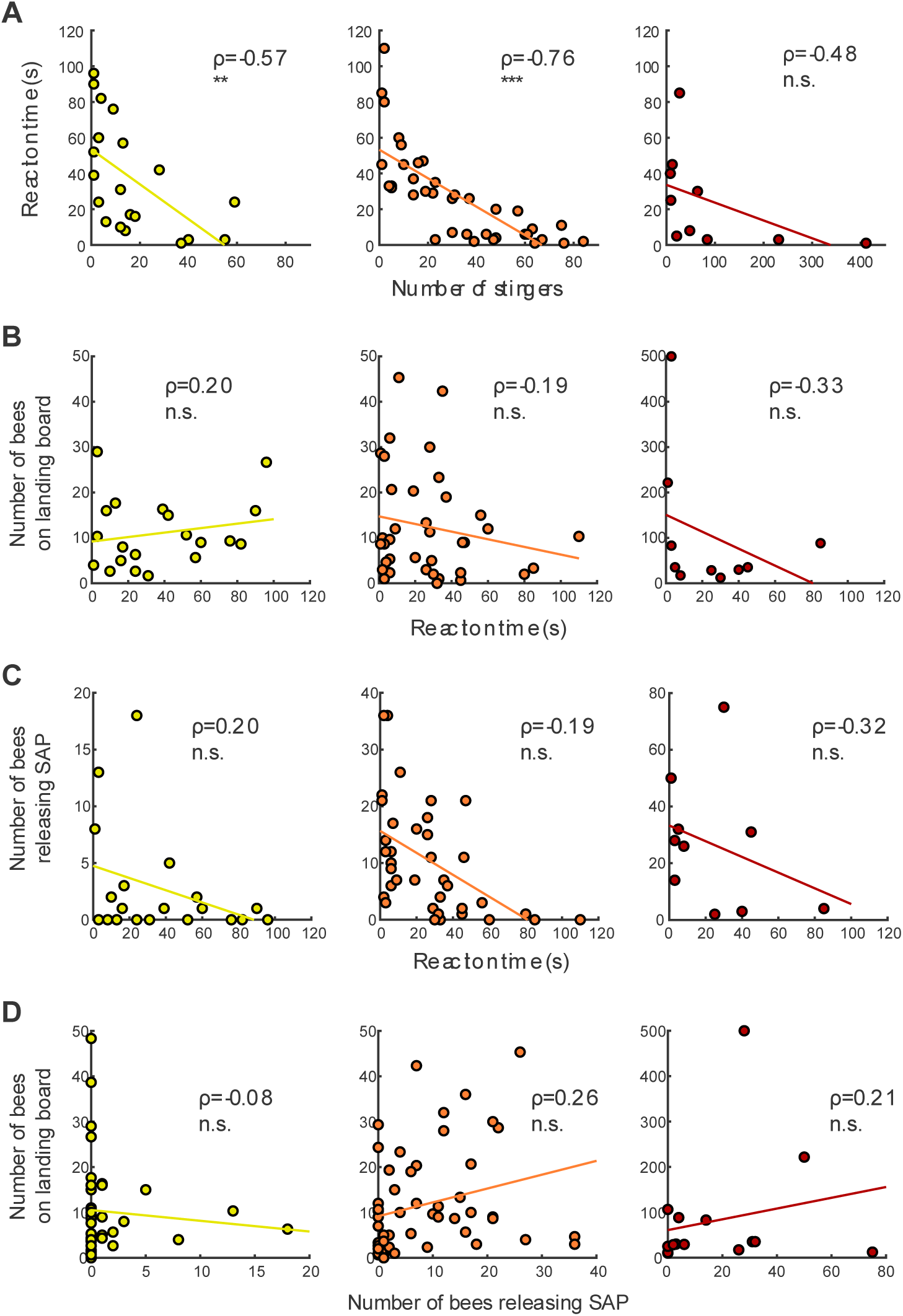
Comparisons between *A. m. carnica*, *A. m. mellifera* and *A. m. scutellata-* continued. A. Correlations between the number of stingers delivered to the flag and the reaction time. B. Correlations between the reaction time and the number of bees on the landing board before the disturbance. C. Correlations between the reaction time and the number of bees actively releasing the SAP during the disturbance. D. Correlations between the number of bees actively releasing the SAP during the disturbance and the number of bees on the landing board before the disturbance. *carnica* (first column, yellow): N=5 colonies, n=38 trials (7-8 per colony); *mellifera* (middle column, orange): N=7 colonies, n=57 (7-9 per colony); *scutellata* (last column, red): N=3 colonies, n=15 trials (3-6 per colony).

**Supplementary Figure 2:**
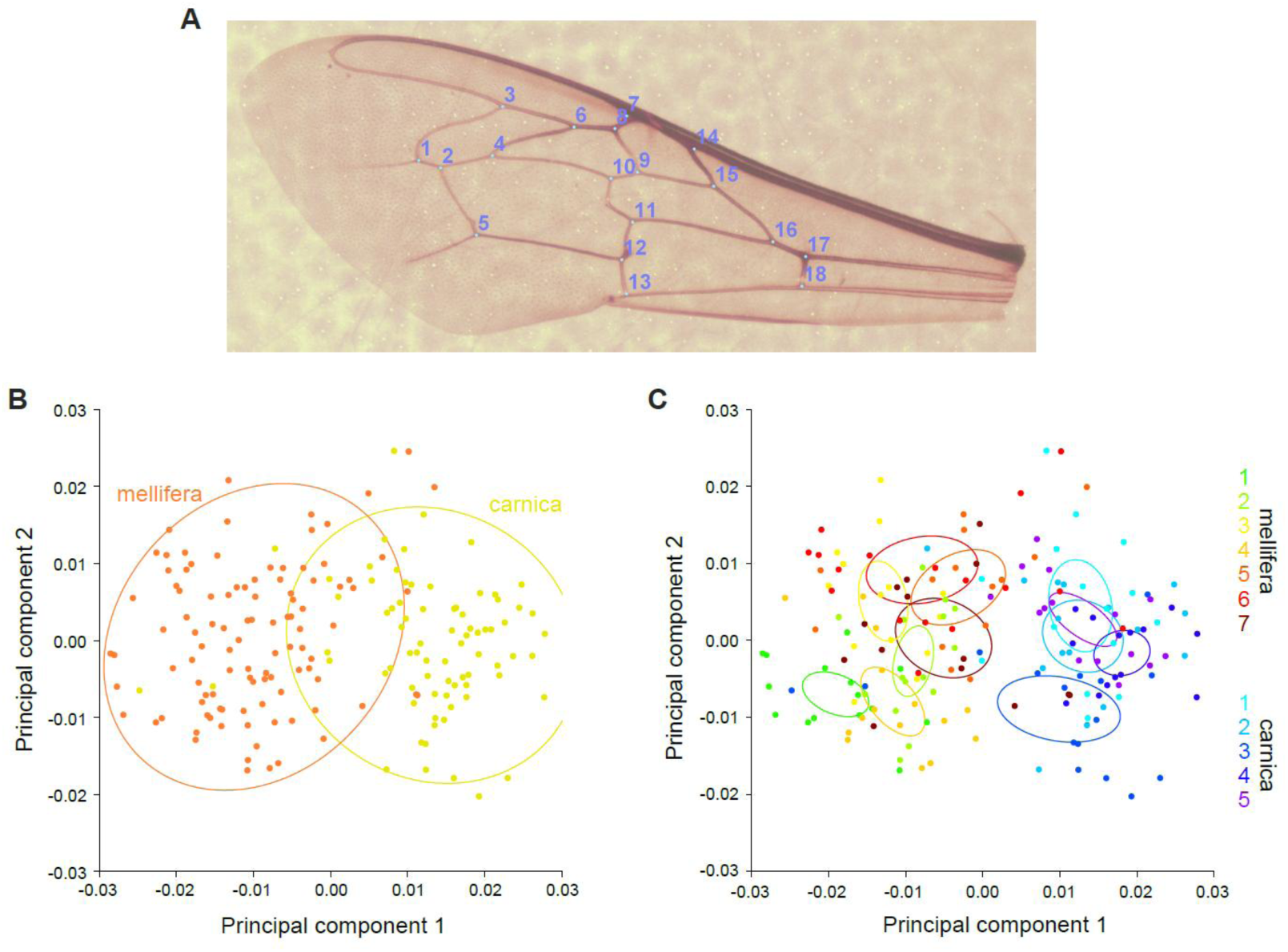
Morphometric analysis of European populations. A. Example wing with annotated landmarks. B. PCA separates individuals in accordance with their subspecies (permutation test: p<0.001). C. Grouping at the colony level on the same principal components also reveals a clear separation between the 2 subspecies. Importantly, no colony occupies an intermediate position (which could have been a sign of hybridization).

**Supplementary Figure 3:**
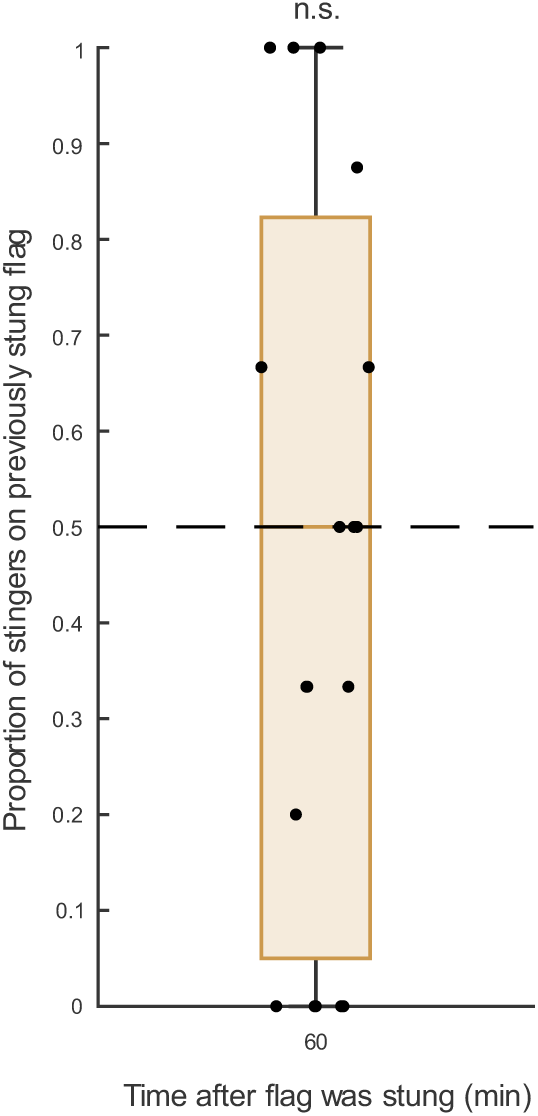
Marking of enemies in European bees, with higher initial stinger load. Proportion of stingers on previously stung flag after 60 min, with an initial load of 33.7 stingers on average. N=6 colonies, n=19 trials.

## References

1. Ruttner, F. (1988). Biogeography and Taxonomy of Honeybees 10.1007/978-3-642-72649-1.

2. Dlugosch, K.M., and Parker, I.M. (2008). Founding events in species invasions: genetic variation, adaptive evolution, and the role of multiple introductions. Mol. Ecol. 17, 431–449. 10.1111/j.1365-294x.2007.03538.x.

3. Colautti, R.I., Alexander, J.M., Dlugosch, K.M., Keller, S.R., and Sultan, S.E. (2017). Invasions and extinctions through the looking glass of evolutionary ecology. Philosophical Transactions of the Royal Society B: Biological Sciences 372. 10.1098/rstb.2016.0031.

4. Collins, A.M., Rinderer, T.E., Harbo, J.R., and Bolten, A.B. (1982). Colony defense by Africanized and European honey bees. Science 218, 72–74.

5. Breed, M.D., Guzman-Novoa, E., and Hunt, G.J. (2004). Defensive behavior of honey bees: organization, genetics, and comparisons with other bees. Annu. Rev. Entomol. 49, 271–298. 10.1146/annurev.ento.49.061802.123155.

6. Guzman-Novoa, E., Hunt, G.J., Uribe-Rubio, J.L., and Prieto-Merlos, D. (2004). Genotypic effects of honey bee (*Apis mellifera*) defensive behavior at the individual and colony levels: the relationship of guarding, pursuing and stinging. Apidologie 35, 15–24. 10.1051/apido:2003061.

7. Rinderer, T.E. (1988). Evolutionary aspects of the Africanization of honeybee populations in the Americas. In Africanized Bees and Bee Mites, G. Needham, R.E.J. Page, M. Delfinado-Baker, and C. Bowman, eds. (Ellis Horwood Ltd), pp. 13–28.

8. Spivak, M., Fletcher, D.J.C., and Breed, M.D. (1991). The “African” honey bee (CRC Press).

9. Alaux, C., Sinha, S., Hasadsri, L., Hunt, G.J., Guzman-Novoa, E., DeGrandi-Hoffman, G., Uribe-Rubio, J.L., Southey, B.R., Rodriguez-Zas, S., and Robinson, G.E. (2009). Honey bee aggression supports a link between gene regulation and behavioral evolution. Proc. Natl. Acad. Sci. USA *106*, 15400-15405. 10.1073/pnas.0907043106.

10. Hunt, G.J. (2007). Flight and fight: a comparative view of the neurophysiology and genetics of honey bee defensive behavior. J. Insect Physiol. 53, 399–410. 10.1016/j.jinsphys.2007.01.010.

11. Fletcher, D.J.C. (1978). The African Bee, *Apis Mellifera Adansonii*, in Africa. Annu. Rev. Entomol. 23, 151–171. 10.1146/annurev.en.23.010178.001055.

12. Schneider, S.S., and MacNally, L.C. (1992). Colony defense in the African honey bee in Africa (Hymenoptera: *Apidae*). Environ. Entomol. 21, 1362–1370.

13. Hepburn, H.R., and Radloff, S.E. (1998). Colony Defense. In Honeybees of Africa, H.R. Hepburn, and S.E. Radloff, eds. (Springer Berlin Heidelberg), pp. 227–241. 10.1007/978-3-662-03604-4_9.

14. Zarate, D., Mukogawa, B., Kohn, J., and Nieh, J.C. (2023). Seasonal variation in defense behavior in European and scutellata-hybrid honey bees (Apis mellifera) in Southern California. Sci Rep 13, 12790. 10.1038/s41598-023-38153-2.

15. Roubik, D.W. (1989). Ecology and natural history of tropical bees (Cambridge University Press). 10.1017/CBO9780511574641.

16. Winston, M.L. (1992). Killer bees. The Africanized honey bee in the Americas (Harvard University Press).

17. Cridland, J.M., Tsutsui, N.D., and Ramírez, S.R. (2017). The Complex Demographic History and Evolutionary Origin of the Western Honey Bee, Apis Mellifera. Genome Biology and Evolution 9, 457-472. 10.1093/gbe/evx009.

18. Dogantzis, K.A., Tiwari, T., Conflitti, I.M., Dey, A., Patch, H.M., Muli, E.M., Garnery, L., Whitfield, C.W., Stolle, E., Alqarni, A.S., et al. (2021). Thrice out of Asia and the adaptive radiation of the western honey bee. Science Advances 7, eabj2151. 10.1126/sciadv.abj2151.

19. Han, F., Wallberg, A., and Webster, M.T. (2012). From where did the Western honeybee (*Apis mellifera*) originate? Ecology and Evolution 2, 1949–1957. 10.1002/ece3.312.

20. Wallberg, A., Han, F., Wellhagen, G., Dahle, B., Kawata, M., Haddad, N., Simões, Z.L.P., Allsopp, M.H., Kandemir, I., De La Rúa, P., et al. (2014). A worldwide survey of genome sequence variation provides insight into the evolutionary history of the honeybee Apis mellifera. Nat. Genet. 46, 1081-1088. 10.1038/ng.3077.

21. Whitfield, C.W., Behura, S.K., Berlocher, S.H., Clark, A.G., Johnston, J.S., Sheppard, W.S., Smith, D.R., Suarez, A.V., Weaver, D., and Tsutsui, N.D. (2006). Thrice out of Africa: ancient and recent expansions of the honey bee, Apis mellifera. Science 314, 642–645. 10.1126/science.1132772.

22. Nouvian, M., Reinhard, J., and Giurfa, M. (2016). The defensive response of the honeybee *Apis mellifera*. J. Exp. Biol. 219, 3505-3517. 10.1242/jeb.143016.

23. Ramirez-Moreno, D., Galizia, C.G., and Nouvian, M. (2025). Division of labour during honeybee colony defence: poetic and scientific views. Philosophical Transactions of the Royal Society B: Biological Sciences 380, 20230272. 10.1098/rstb.2023.0272.

24. Collins, A.M., Rinderer, T.E., Tucker, K.W., and Pesante, D.G. (1987). Response to alarm pheromone by European and Africanized honeybees. J. Apic. Res. 26, 217-223.

25. Breed, M.D., Robinson, G.E., and Page, R.E. (1990). Division of labor during honey bee colony defense. Behav. Ecol. Sociobiol. 27, 395–401.

26. Chwalibog, A. (2017). The Honey Badger in South Africa: Biology and Conservation. International International Journal of Avian & Wildlife Biology 2. 10.15406/ijawb.2017.02.00016.

27. Vanderhaar, J.M., and Hwang, Y.T. (2003). Mellivora capensis. Mammalian Species 721, 1–8. 10.1644/721.

28. Begg, K.S. (2001). Report on the conflict between beekeepers and honey badgers *Mellivora capensis*, with reference to their conservation status and distribution in South Africa. Unpublished report for the Endangered Wildlife Trust, Johannesburg.

29. Hunt, G.J., Wood, K.V., Guzmán-Novoa, E., Lee, H.D., Rothwell, A.P., and Bonham, C.C. (2003). Discovery of 3-Methyl-2-Buten-1-Yl Acetate, a New Alarm Component in the Sting Apparatus of Africanized Honeybees. J. Chem. Ecol. 29, 453-463. 10.1023/a:1022694330868.

30. Wager, B.R., and Breed, M.D. (2000). Does honey bee sting alarm pheromone give orientation information to defensive bees? Ann. Entomol. Soc. Am. 93, 1329–1332. 10.1603/0013-8746(2000)093[1329:DHBSAP]2.0.CO;2.

31. Collins, A.M., Rinderer, T.E., Daly, H.V., Harbo, J.R., and Pesante, D. (1989). Alarm pheromone production by two honeybee (Apis mellifera) types. J. Chem. Ecol. 15, 1747–1756. 10.1007/bf01012262.

32. Nelson, D.R., Collins, A.M., Hellmich, R.L., Jones, R.T., Helm, R.M., Squillace, D.L., and Yunginger, J.W. (1990). Biochemical and immunochemical comparison of Africanized and European honeybee venoms. J. Allergy Clin. Immunol. 85, 80–85. 10.1016/0091-6749(90)90225-s.

33. Resende, V.M.F., Vasilj, A., Santos, K.S., Palma, M.S., and Shevchenko, A. (2013). Proteome and phosphoproteome of Africanized and European honeybee venoms. Proteomics 13, 2638–2648. 10.1002/pmic.201300038.

34. Schumacher, M.J., Schmidt, J.O., Egen, N.B., and Lowry, J.E. (1990). Quantity, Analysis, and Lethality of European and Africanized Honey Bee Venoms. The American Journal of Tropical Medicine and Hygiene 43, 79–86. 10.4269/ajtmh.1990.43.79.

35. Henriques, D., Browne, K.A., Barnett, M.W., Parejo, M., Kryger, P., Freeman, T.C., Muñoz, I., Garnery, L., Highet, F., Jonhston, J.S., et al. (2018). High sample throughput genotyping for estimating C-lineage introgression in the dark honeybee: an accurate and cost-effective SNP-based tool. Scientific Reports 8. 10.1038/s41598-018-26932-1.

36. Tofilski, A. (2008). Using geometric morphometrics and standard morphometry to discriminate three honeybee subspecies. Apidologie 39, 558-563. 10.1051/apido:2008037.

37. Rohlf, F.J. (2021). TpsDig.

38. Nielsen, D.I., Ebert, P.R., Hunt, G.J., Guzmán-Novoa, E., Kinnee, S.A., and Page, R.E. (1999). Identification of Africanized Honey Bees (Hymenoptera: Apidae) Incorporating Morphometrics and an Improved Polymerase Chain Reaction Mitotyping Procedure. Ann. Entomol. Soc. Am. 92, 167–174. 10.1093/aesa/92.2.167.

39. Benjamini, Y., and Hochberg, Y. (1995). Controlling the False Discovery Rate: A Practical and Powerful Approach to Multiple Testing. Journal of the Royal Statistical Society: Series B (Methodological) 57, 289–300. 10.1111/j.2517-6161.1995.tb02031.x.

